# Impacts of thermal fluctuations on heat tolerance and its metabolomic basis across plant and animal species

**DOI:** 10.1101/2020.07.23.217323

**Authors:** Natasja Krog Noer, Majken Pagter, Simon Bahrndorff, Anders Malmendal, Torsten Nygaard Kristensen

## Abstract

Temperature varies on a daily and seasonal scale and thermal fluctuations are likely to become even more pronounced under future climate changes. Studies suggest that plastic responses are crucial for species’ ability to cope with thermal stress, but traditionally laboratory studies on ectotherms are performed at constant temperatures and often limited to a few model species and thus not representative for the natural environment. We argue that thermoregulatory behavior and microhabitat shape the response exerted by different organisms to fluctuating temperatures. Thus, a sessile organism incapable of significant behavioral temperature avoidance will be more plastic and exert greater physiological response to thermal fluctuations than mobile organisms that can quickly evade temperature stress. Here we investigate how acclimation to fluctuating (13.2-26.9°C) and constant (20.4°C) temperatures impact heat stress tolerance across a plant (*Arabidopsis thaliana*) and two animal species (*Orchesella cincta* and *Drosophila melanogaster*) inhabiting widely different thermal microhabitats and selective pasts. Moreover, we investigate the underlying metabolic responses of acclimation using an NMR metabolomics approach. We find increased heat tolerance for all species exposed to fluctuating acclimation temperatures; most pronounced for *A. thaliana* which also showed a strong metabolic response to thermal fluctuations. Generally, sugars were more abundant across *A. thaliana* and *D. melanogaster* when exposed to fluctuating compared to constant temperatures, whereas amino acids were less abundant. However, we do not find much evidence for similar metabolomics responses to fluctuating temperature acclimation across species. Differences between the investigated species’ ecology, their distinct selective past and different ability to behaviorally thermoregulate may have shaped their physiological response to thermal fluctuations.

## 1. Introduction

The natural environment is constantly changing and abiotic factors fluctuate continuously on various spatial and temporal scales. These changes can pose stress on organisms [1,2] and especially extreme temperatures are important physical factors that affect the abundance and distribution of species [3-7]. To survive and reproduce in fluctuating and periodically stressful environments individuals and populations need to adjust their physiology, morphology or behavior to mitigate adverse effects on fitness. This can occur via phenotypic changes within the lifetime of an individual, cross-generational epigenetic responses, or evolutionary adaptations in populations across generations [4,8]. Most laboratory studies on physiological and evolutionary adaptation to thermal stress on ectotherms have used constant thermal conditions to address responses and impacts [2,9]. However, under natural conditions temperatures are rarely constant and even less so in the face of climate change causing increased variability and decreased predictability of temperatures [10,11]. Recent studies have shown that thermal variability can have significant impact on thermal reaction norms and that conclusions drawn on the basis of studies in constant temperature environments does not always hold under variable thermal conditions [12-15]. For instance, several studies on insect species have reported that acclimation to thermal fluctuations will benefit heat tolerance due to Jensen’s inequality, i.e. the time spent closer to optimum temperatures exceeds the time spent in less favorable conditions when temperature fluctuates [16-18]. However, these findings have been inconsistent and likely depend on species- and trait-specific levels of plasticity and the nature of the thermal regime [19,20].

A common expectation for studies examining the effect of thermal variability on heat and cold stress resistance is that environmental heterogeneity and variability in habitat temperatures will select for high levels of plasticity in thermal tolerance traits compared to stable thermal environments [21-26]. Under the current climate change projections, which predict fast changes in e.g. temperature within days or seasons, plasticity will thus be important for survival when there is little time for evolutionary responses [27-32]. Many studies have examined if associations exists between environmental temperature variability of natural habitats and the degree of plasticity of thermal tolerance in animal ectotherms (reviewed by Addo-Bediako et al. [21]; Hoffmann et al. [30]; see also[33-35]), and plants [36-39]. In the studies with animals no relationship between the latitudinal origin of populations and the level of plasticity in thermotolerance is found, whereas the findings are conflicting for plant studies. However, local differences in plasticity levels are often observed (at similar latitudes populations may vary in strategies used to cope with thermal challenges) suggesting that populations are adapted to local microhabitats which disrupt coarse-scale clines in climatic variables and also permits behavioral thermoregulation [7]. Thermoregulatory behavior reduces the temperature variation experienced by some invertebrates because they actively seek shade, bask in the sun, or move between sun and shade [40,41]. However, the capacity for behavioral thermoregulation will depend on the mobility of the species [42]. Contrarily, plants have limited capability of thermoregulatory behavior on a short time scale (e.g. daily) and this can explain why plant studies generally find stronger evidence for increased plasticity levels with latitude compared to studies on ectotherms. The sessile life strategy might thus promote selection on alternative physiological, biochemical, or morphological mechanisms of stress avoidance and mitigation [28,37,40,43].

Metabolomics has been recognized as a valuable tool for understanding the mechanisms underlying acclimation processes in plants and animals as it provides an integrated measure of regulatory processes at the different molecular levels, combined with external environmental influences [44,45]. All of these processes are reflected in the metabolome, which is closely linked to the observed functional phenotype. Thus, the metabolome might constitute a reliable predictor of organismal phenotypes and provide novel insight into the underpinnings of complex traits such as responses to thermal fluctuations [46,47] Here, we investigate tolerance to heat stress in the hexapods *Drosophila melanogaster and Orchesella cincta*, and the plant *Arabidopsis thaliana*. We acclimated the three species at either constant or fluctuating temperatures, but with the same mean temperature. We expect to find that individuals exposed to thermal fluctuations will have higher heat tolerance compared to those exposed to constant thermal acclimation conditions prior to testing. We further investigate the metabolic consequences of exposure to respectively constant and fluctuating temperatures using NMR metabolomics hypothesizing that both shared and distinct responses to thermal fluctuations are observed across the three species.

## 2. Materials and Methods

### 2.1. Species and populations used for experiments

The experiment was performed in two independent experimental runs with sampling for NMR metabolomics in the second run.

#### 2.1.1. Drosophila melanogaster

Flies used in the study were from a population of *D. melanogaster* that was set up in 2010 using the offspring of 589 inseminated females caught at Karensminde fruit farm in Odder, Denmark. The population was maintained in the laboratory for ca. 220 generations in the lab at a population size >1000 individuals prior to performing the experiments reported here (for further details see: Schou et al. [48]). The flies were held in plastic bottles containing 50 mL agar-sugar-yeast-oatmeal standard Drosophila medium [49] at a density of approximately 200 flies per bottles and maintained at 19°C in a 12:12 h light/dark regime. The experimental flies were produced by density-controlled egg laying in 10 bottles of 200 flies for 6 hours. Newly eclosed flies were transferred to bottles with fresh media within 12 hours of eclosion. One day prior to acclimation start the flies were sexed under light CO_2_ anesthesia (<5 min) and groups of 10-12 males were placed in 35 mL vials containing 7 mL standard Drosophila medium. Flies were 5-6 days of age at acclimation start for both experimental runs. During acclimation, fresh media was provided every second day.

#### 2.1.2. Orchesella cincta

Collembolans used in the experiment originated from a population collected in Siena, Italy, in 2016 and thereafter maintained at 20°C, 70% RH and a photoperiod of 12:12 hour light:dark regime for ca. 15 generations (for details see Jensen et al., 2019). During acclimation, collembolans were held in 55 mm petri dishes containing a water-saturated plaster-of-paris:charcoal (9:1) medium and an algae-covered twig was provided as food source. Each petri dish contained 10-13 individuals that were 8 weeks old at acclimation start in the first experimental run and 10 weeks old at the second run. A few drops of water were added to each petri dish every day to prevent desiccation stress and fresh food was provided every second day.

#### 2.1.3. Arabidopsis thaliana

*A. thaliana* seeds used in the study were from the Columbia-0 (Col-0) accession. Seeds were surface sterilized in 70% (v/v) ethanol for 10 min and then briefly mixed with 100% (v/v) ethanol before drying on a piece of sterile filter paper. Surface-sterilized seeds were stratified in sterile water at 4°C for three days prior to plating on 55 mm petri dishes containing 1x Murashige and Skoog (MS) basal growth medium (Duchefa), 0.5 gL^−1^ MES and 1.0% (w/v) agar at pH 5.7. Seeds were plated at a density of 24 seeds per plate. Plates were sealed with Micropore tape and placed horizontally in a growth room at 20°C with an 8 h day length at 150 µmol m^−2^ s^−1^ for three days to allow seedlings to emerge, after which the acclimation treatments were initiated.

#### 2.2. Thermal Acclimation Regimes

A constant and a fluctuating thermal regime with equal mean temperature and a photoperiod of 8:16 h light/dark and light intensity of 150 µmol m^−2^ s^−1^ were generated in two programmable Plant Growth Chambers (Snijders Microclima MC1750E). The constant regime retained a temperature of 20.4± 0.2°C throughout the day and night. The fluctuating thermal regime varied predictably and diurnally around the mean temperature 20.4°C, reaching a minimum temperature of 13.2± 0.1°C, at rate of 0.04°C min^−1^, early in the morning, and a maximum temperature of 26.9± 0.1°C, at a rate of 0.06°C min^−1^, in the afternoon (see Fig. 1).

**Figure 1:**
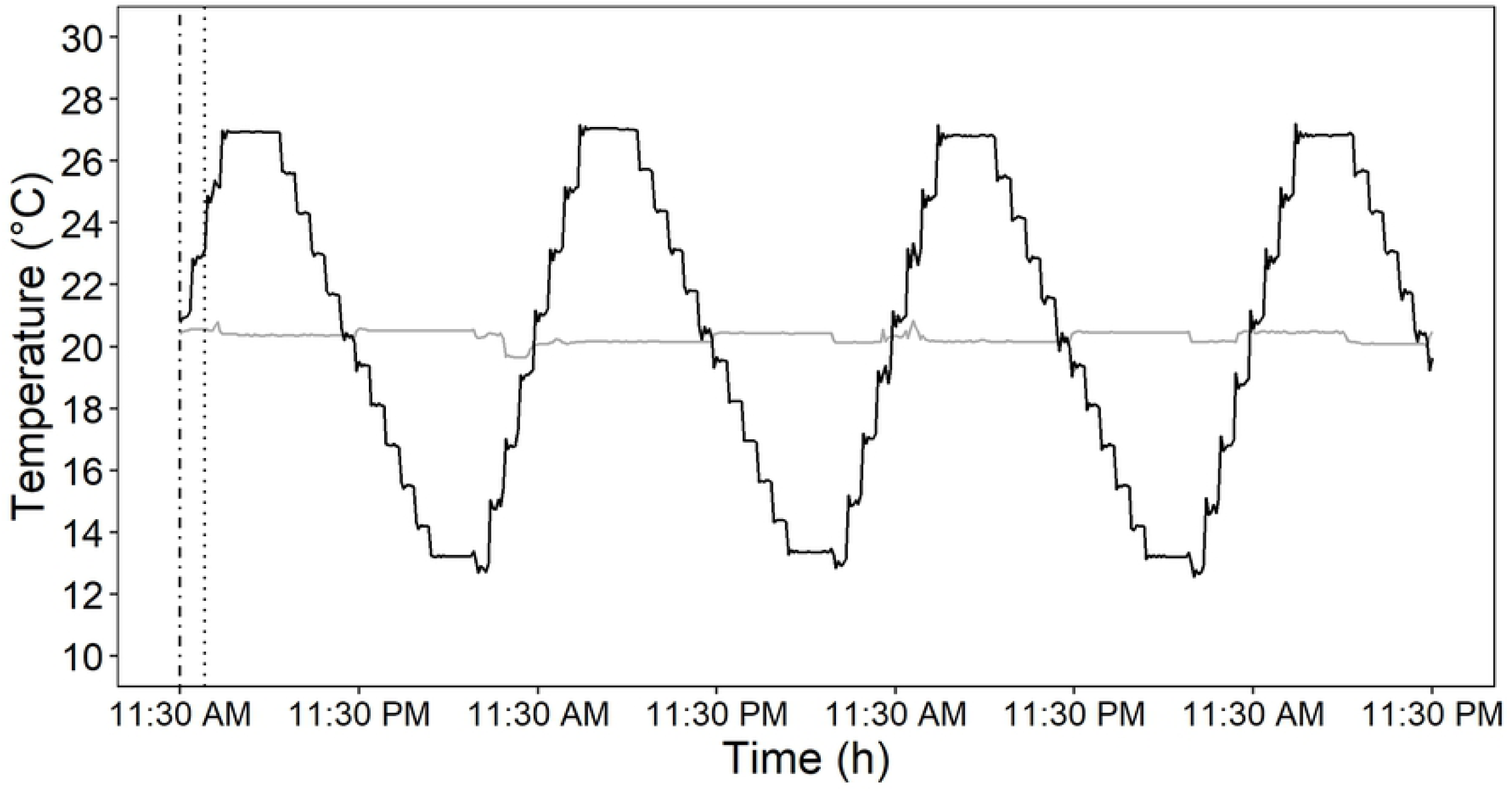
Section of the recorded temperatures in the constant (grey) and fluctuating (black) thermal regimes from the second experimental run. The constant thermal regime maintained a temperature of 20.4± 0.2°C and the fluctuating regime cycled around the mean temperature in intervals of 13.2-26.9°C. Adult flies and collembolans and plant seedlings were acclimated to each acclimation regime for 6-7 days. The experiment was performed twice and the dotted and dashed lines represent the temperature at the initiation of acclimation treatments in the first and second experimental replicate, respectively.

Acclimation treatments were started by relocating 90 (replicate) petri dishes containing three-day old *A. thaliana* seedlings or *O. cincta*, and 90 Drosophila vials containing *D. melanogaster*, randomly to each acclimation treatment. The temperatures in the constant and fluctuating regimes were 20.4 and 24.3°C at acclimation start for all organisms in the first experimental run, and 20.4°C in both regimes in the second run (Fig. 1). The replicate petri dishes and vials were reshuffled in a randomized manner inside the chambers once a day during the acclimation period to minimize the internal differences in temperatures and light that individual replicates were exposed to. Flies and collembolans were acclimated for 7 days, and seedlings for 6 days before assessment of their heat tolerance.

### 2.3. Heat tolerance

Heat tolerance was tested using a heat mortality assay exposing the organisms to 7-8 species specific temperatures ranging from non-lethal to lethal: *D. melanogaster* were exposed to 35, 36, 37, 38, 39, 40, 41°C; *O. cincta* to 35, 36, 37, 38, 39, 40, 41, 42°C; *A. thaliana* to 37, 39, 41, 43, 44, 45, 47, 49°C. The species-specific test temperatures were based on pilot tests (results not shown). Thermal incubators were used to generate the different test temperatures and the tests were conducted at midday when the temperature in the two thermal regimes was the same; ∼20.4°C. This was done to minimize effects of different temperatures and daily rhythm on heat tolerance. For flies and collembolans, 10 replicates of 10 individuals from each acclimation treatment were exposed to each stress temperature for 1 hour. During the tests the flies were kept in plastic vials containing 7 mL standard Drosophila medium and collembolans were kept on Plaster-of-Paris medium. Following exposure to heat stress, *D. melanogaster* and *O. cincta* recovered for 1 hour at 20.4°C and subsequently the mortality was scored as the number of dead individuals out of the total number of individuals in each replicate. For *A. thaliana*, 10 replicate plates with ∼24 seedlings from each acclimation treatment were likewise exposed to each of the chosen test temperatures for 1 hour and then returned to the growth chamber at a constant temperature of 20.4°C. The seedlings were kept in the petri dishes containing (MS) medium during the test and recovery. Thermal damages in plants build up slowly when exposed to moderately high temperatures [43] and to ensure that we had accounted for the total mortality caused by thermal stress the number of viable seedlings was quantified after both 4 and 7 days of recovery. Seedlings that were still green and produced new leaves were scored as survivors.

### 2.4. NMR

In the second experimental run, hexapod and plant samples were snap-frozen in liquid nitrogen for metabolome analysis while other individuals were tested for heat tolerance simultaneously. Metabolites were extracted from 6 replicates of 10 male flies and 6 replicates of 10 non-sexed collembolans from each acclimation treatment using the same protocol as described by Ørsted et al. [50] and from 5 replicates of 24 seedlings from each acclimation treatment for *A. thaliana* using an extended protocol. Whole-body tissues from each sample were mechanically homogenized in 1 mL of Acetonitrile solution (50%, 50% ddH_2_O) using sterile glass beads and a homogenizer (FastPrep-24™ -MP Biomedicals) for 2 x 35 sec at 3800 rpm. Plant samples were sonicated for 15 minutes at room temperature prior to the proceeding steps. All samples were cooled on ice, centrifuged at 14,000 rpm for 10 min at 4°C. The supernatant was transferred to new tubes, snap frozen, lyophilized, and stored at -80°C until NMR analysis.

NMR measurements were performed at 25°C on a Bruker Avance III HD 800 spectrometer (Bruker Biospin, Rheinstetten, Germany), operating at a ^1^H frequency of 799.87 MHz, and equipped with a 3 mm TCI cold probe. ^1^H NMR spectra were acquired using a standard NOESYPR1D experiment with a 100 ms delay. A total of 128 transients of 32 K data points spanning a spectral width of 20 ppm were collected. The spectra were processed using Topspin (Bruker Biospin, Rheinstetten, Germany). An exponential line-broadening of 0.3 Hz was applied to the free-induction decay prior to Fourier transformation. All spectra were referenced to the DSS signal at 0 ppm, manually phased and baseline corrected. The spectra were aligned using icoshift [51]. The region around the residual water signal (4.87-4. 70 ppm) was removed in order for the water signal not to interfere with the analysis. The high- and low-field ends of the spectrum, where no signals except the reference signal from DSS appear, were also removed (i.e., leaving data between 9.7 and 0.7 ppm).

### 2.5. Data analysis

#### 2.5.1. Thermal tolerance

We fitted a generalized linear model with a binomial link function on survival proportions for each species. This allowed us to look for batch effects between repeated experiments. A likelihood ratio test (LRT) was used to compare a model containing an interaction term between temperature and experimental run with a reduced model omitting this term. All LRTs were significant which indicated significant effects of experimental run and the data was treated as two independent experiments for the rest of the analysis.

For every heat exposure temperature, survival was calculated as the number of survivors over total number of individuals for each replicate. The Lethal median Temperature (LT_50_) for each acclimation treatment was found by logistic regression on survival proportions for each stress temperature using the drc-package in R [52]. Significant differences in LT_50_ for each species were found by comparing confidence intervals of LT_50_ estimates and by Chi-square LRT on a model incorporating differences in LT_50_ between the two thermal regimes and a model assuming common LT_50_ for both.

#### 2.5.2. NMR data

Multivariate analyses were performed on spectral data that was normalized by probabilistic quotient area normalization [53] to suppress separation caused by variation in sample volumes, and pareto-scaled to reduce variance caused by metabolite differences. Principal component analysis (PCA) was used to differentiate metabolite content between species and thermal acclimation treatment. Significant effects of species and acclimation regime were tested on PCA scores using MANOVA.

Metabolome differences caused by thermal acclimation regime were further assessed using Orthogonal Projection to Latent Structures Discriminant Analysis (OPLS-DA) on the data combined for all species and for individual species. Validation scores for the OPLS-DA models were calculated by 7-fold cross-validation. The analysis is regression-based and seeks to correlate sample variation with a response vector that contains sample information (acclimation treatment) while finding uncorrelated variation (orthogonal components) that are systematic in the data. This analysis is useful when the effects of interest are masked by variables that have larger influences on the sample variation, e.g. species metabolome variation [54,55]. OPLS-DA was carried out using the SIMCA16 software (Umetrics, Malmö, Sweden).

In order to identify the significant changes in metabolite concentrations when going from constant to fluctuating acclimation regimes the OPLS-DA loadings (amplitude and correlation) were plotted for all models. The correlations were calculated after removing the variation explained by the orthogonal components. The threshold for significant change in metabolites between acclimation treatments were *P* < 0.05 after correction for multiple testing for 50 metabolites. Relative changes in metabolite concentrations in organisms acclimated to the two thermal regimes were calculated for every significant metabolite as the difference in concentration in individuals from the constant and the fluctuating acclimation treatments divided by the median concentration in individuals from the constant acclimation regime.

## 3. Results

### 3.1. Thermal fluctuations increase heat tolerance

Acclimation to a fluctuating thermal regime compared to a constant regime with equal mean temperature consequently increased LT_50_ for all species in both experimental runs (Table 1, suppl. Fig. S1). The greatest difference was observed for *A. thaliana* where LT_50_ for plants exposed to the fluctuating thermal acclimation was 0.5°C higher in experimental run 1 and 0.9°C higher in run 2 (Table 1). For *D. melanogaster* these numbers were 0.3 and 0.2°C in run 1 and 2, respectively. An increase in LT_50_ was also observed in *O. cincta* where individuals from the fluctuating acclimation treatment had 0.3 and 0.1°C higher LT_50_ in run 1 and 2, respectively, but only the response in the first experimental run was significant (Table 1). Generally, we observed higher heat tolerance for *A. thaliana* than for both invertebrates, whereas only slightly higher LT_50_ was observed in *O. cincta* compared to *D. melanogaster*.

**Table 1:** LT_50_ values (mean ± SE) for species acclimated to constant and fluctuating thermal regimes. Significant differences in LT_50_ for the two acclimation treatments were found by chi-square test for each species.

| Species | Acclimation treatment |  | Chi-square test |  |
| --- | --- | --- | --- | --- |
| | Constant | Fluctuating | *p*-value | $\chi^2_{(d.f.)}$ |
| LT50 (°C) |  |  |  |  |
| Experimental run 1 |  |  |  |  |
| *Drosophila melanogaster* | 38.27 (±0.07) | 38.56 (±0.07) | 0.002 | 9.42<sub>(1)</sub> |
| *Orchesella cincta* | 39.69 (±0.09) | 39.94 (±0.08) | 0.034 | 4.49<sub>(1)</sub> |
| *Arabidopsis thaliana* | 44.79 (±0.05) | 45.33 (±0.07) | < .001 | 53.71<sub>(1)</sub> |
| Experimental run 2 |  |  |  |  |
| *Drosophila melanogaster* | 38.82 (±0.06) | 39.01 (±0.06) | 0.028 | 4.81<sub>(1)</sub> |
| *Orchesella cincta* | 39.30 (±0.08) | 39.38 (±0.08) | 0.460 | 0.55<sub>(1)</sub> |
| *Arabidopsis thaliana* | 43.75 (±0.05) | 44.64 (±0.05) | < .001 | 150.16<sub>(1)</sub> |

### 3.2. Effect of thermal fluctuations on the metabolome

Principal component analysis (PCA) was performed on the combined and separate metabolite spectra of *D. melanogaster, O. cincta*, and *A. thaliana* to characterize the metabolite response underlying acclimation to constant and fluctuating thermal regimes for all species. The total variation between samples was explained by 33 principal components, however PC1 and PC2 accounted for most of this variation (64.3%, Fig. 2 and S2). An inspection of the PCA scores plotted for PC1 and PC2 shows a distinct separation of metabolites associated with species and this was substantiated by a test for differential metabolite response on PCA scores (MANOVA; *P* < .001). PC1 further separates clusters associated with thermal regime (Fig. 2 and 3), but this effect was not significant (MANOVA; *P* = 0.113).

**Figure 2:**
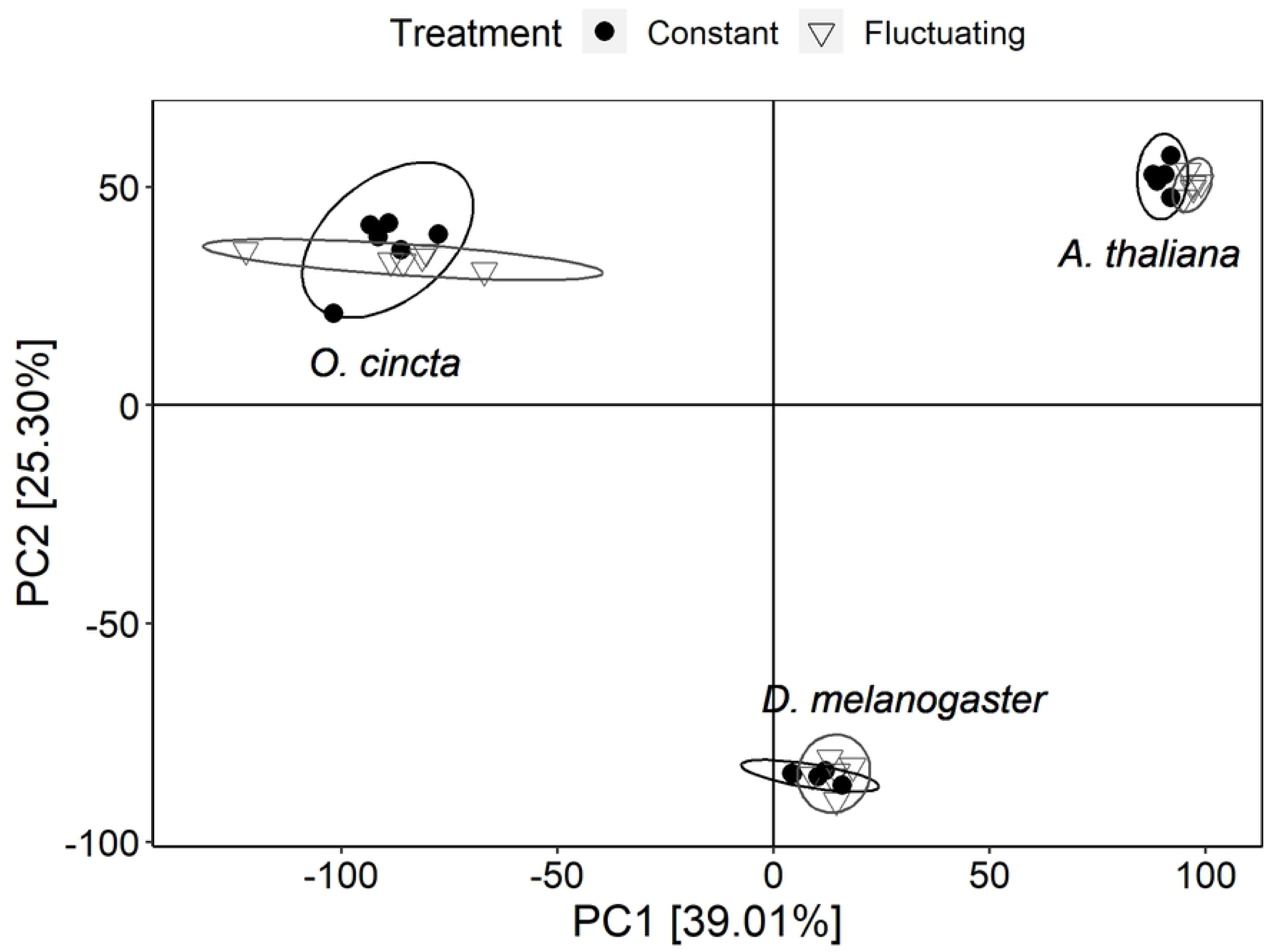
PCA scores plot on metabolites from whole-body extract of *Drosophila melanogaster, Orchesella cincta*, and *Arabidopsis thaliana* acclimated to constant (filled circles) and fluctuating (open triangles) thermal regimes. PC1 and PC2 account for 39.01% and 25.39% of the variance between samples, respectively.

**Figure 3:**
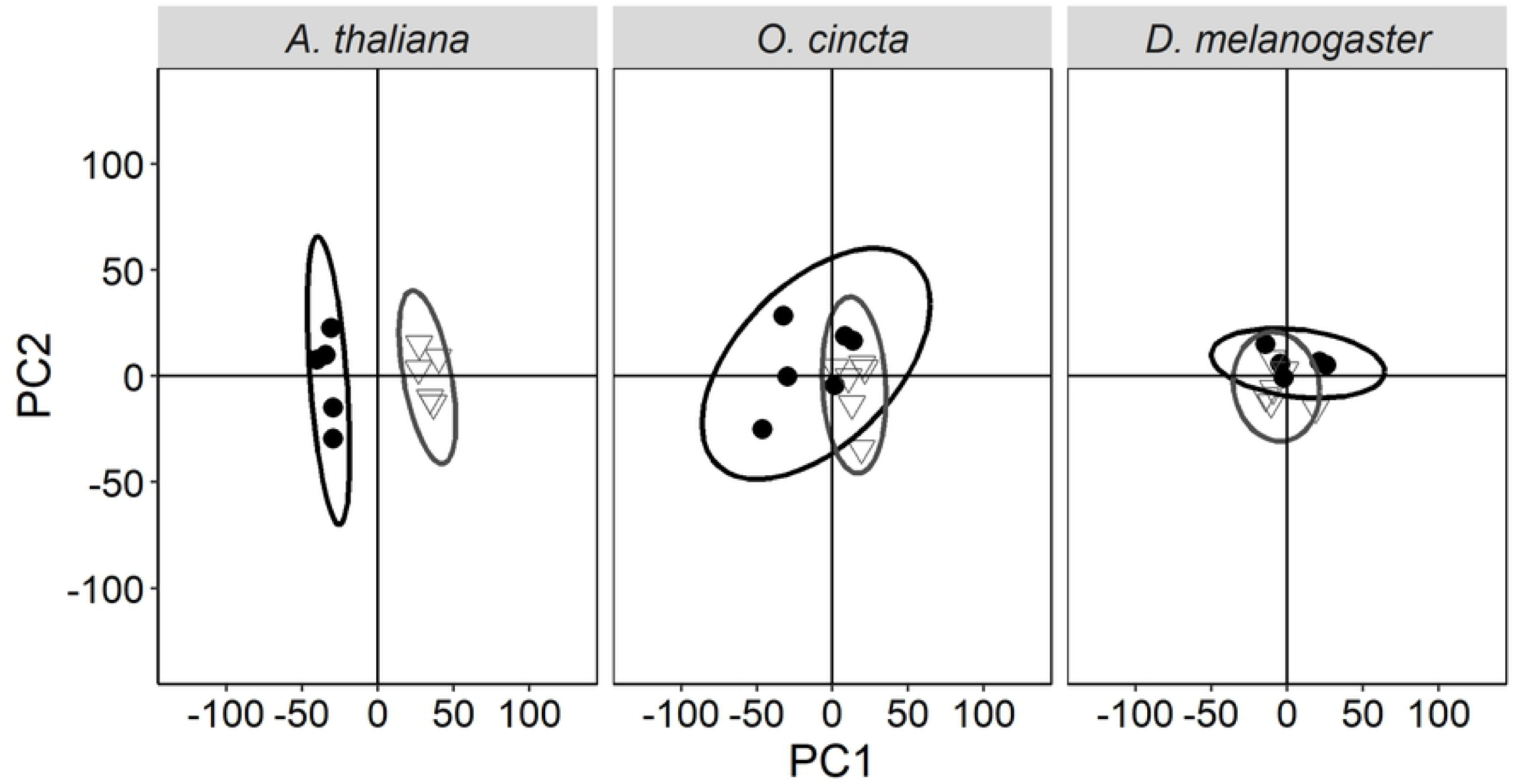
PCA scores plot for *Arabidopsis thaliana, Orchesella cincta, and Drosophila melanogaster* acclimated to constant (filled circles) or fluctuating (open triangles) thermal regimes. Ellipses represent 95% CI.

Because of large variation caused by species-specific metabolite differences, an OPLS-DA was performed to focus the analysis towards differences in acclimation treatment while diminishing variation caused by species. The OPLS-DA model was composed of one predictive component and three orthogonal components (Table S1). Thermal regime accounted for merely 3% of the total metabolite variation in the samples, but the predictive ability of the model to correctly group a sample into constant or fluctuating acclimation treatment based on the metabolite content in the sample was significant (predictability Q^2^Y = 0.5, Table S1).

In addition, OPLS-DA models were performed on individual species (Table S1). All models were of good quality and showed a high correlation between metabolite content and thermal regime. For *A. thaliana* 60% of the total metabolite variation was explained by the acclimation treatment, while the corresponding number was 18% for both *D. melanogaster* and *O. cincta* (Table S1).

The OPLS-DA loadings from each individual OPLS-DA model was used to identify metabolites that differed significantly between individuals acclimated to fluctuating and constant thermal regimes (Fig. 4 and S3). The analysis showed that the set of metabolites that was up- or downregulated differed for each species. Metabolite changes that were significantly associated with the predictive component for *A. thaliana* included elevated levels of glucose and suppressed levels of glutamine (gln), arginine (arg), and gamma-aminobutyric acid (GABA) (Fig. 4 and Fig. S2D). Sucrose levels were elevated in *D. melanogaster* acclimated to thermal fluctuations and alanine (Ala) levels lowered (Fig. 4 and S2C). Lastly, *O. cincta* that was acclimated to thermal fluctuations had elevated levels of hydroxyphenyl ethanol and 3-hydroxybutyric acid (Fig. 4 and S2B).

**Figure 4:**
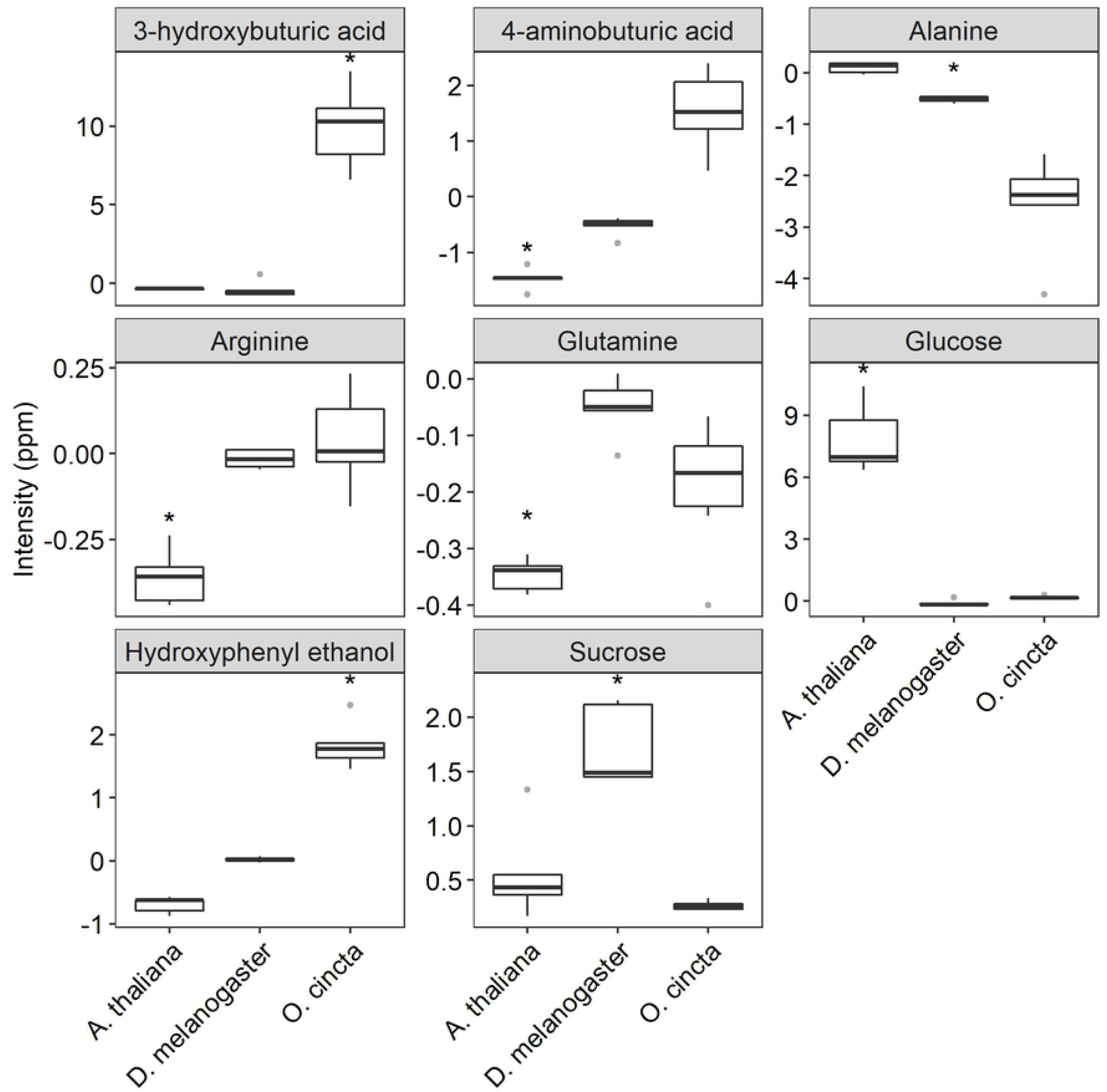
Whisker box plots of change in relative concentrations of nine metabolites in *A. thaliana, D. melanogaster*, and *O. cincta* acclimated to fluctuating and constant temperatures. Significant changes in concentrations are indicated by * (*p < 0.05*). The y-axis shows the concentration of each metabolite in the organisms acclimated to fluctuating temperatures (measured as spectral intensity) relative to the average concentration of the metabolites in organisms acclimated to constant temperatures. The centerline in each box represents the median, and the upper and lower boundaries are the 25^th^ and 75^th^ quantiles. The whiskers mark the extremes, and dots represent the outliers.

## 4. Discussion

In this study we investigated the effect of thermal fluctuations on heat tolerance and the metabolome in three taxonomically distant species. We found that individuals from all three species acclimated at fluctuating temperatures were more heat tolerant than individuals acclimated to constant temperatures (Table 1, Fig. S1). The increased heat tolerance of individuals from the fluctuating temperature regime is in line with previous studies on invertebrate species exposed to predictable thermal fluctuations [16-18], but has to our knowledge not previously been shown for plants species.

Of the three species that we investigated we observed the most pronounced increase in heat tolerance in response to thermal fluctuations for *A. thaliana* (Table 1, Fig. S1). This finding suggests that *A. thaliana* has a larger capacity to adjust its phenotype to environmental fluctuations than *D. melanogaster* and *O. cincta*. This agrees with studies on plants that have found strong correlations between latitudinal temperature variability and plasticity of various fitness-related traits, including photosynthetic rate, water-use efficiency, seed-output, leaf angles and number of flowers [37,56,57]. Such association between latitudinal temperature variability and fitness-related traits and heat tolerance has generally been much weaker for insects [14,15,34,58]. Although we cannot make any conclusions on why plants in general respond stronger to thermal fluctuation based on our study, it could be speculated that plants have developed stronger plastic responses as a consequence of their sessile lifestyle compared to invertebrates. We only found marginal differences in the acclimation response to constant and fluctuating temperatures of flies and collembolans and no significant effect of acclimation treatment on the metabolomes (Table 1 and Fig. 3). This suggests a lower ability of these species to induce a thermal plastic response which may relate to the ability of the species’ to behaviorally evade stressors in nature by e.g. seeking deeper into the soil column or escaping rapidly by flight to more favorable conditions [33,59,60].

The magnitude of the acclimation response observed in LT_50_ values was manifested in the metabolome showing bigger differentiation for *A. thaliana* exposed to fluctuating and constant temperatures compared to the arthropod species investigated (Fig. 2 and 3 and Table S1). This is well in accordance with the idea that plants, due to their limited mobility, exert a greater metabolite response to environmental fluctuations than invertebrates. Further, we found that the set of metabolites that was elevated or suppressed in response to thermal fluctuations for each species differed, but some of the affected metabolites shared some biochemical properties. For instance, sugar levels were elevated in plants and flies exposed to thermal fluctuations, whereas amino acids were suppressed (Fig. 4). These patterns were not found for collembolans, which showed a markedly different metabolite response than flies and plants.

Accumulation of soluble sugars, which act as compatible compounds that help stabilizing proteins and membranes and regulating osmotic pressure, is a common low temperature response in invertebrates [13,33,59-62] and a low and high temperature response in plants [43,63-65]. In our experiment, heat tolerance was tested at midday when temperatures in the fluctuating thermal regime had returned to initial mean temperature subsequent to a cool thermal peak reaching 13°C during the night. It has previously been found that *D. melanogaster* exposed to gradual cooling shows increasing levels of sugars and decreasing levels of amino acid when temperatures approximate 10°C and that the levels remain elevated for up to 4 hours after returning to pre-cooling temperatures [62]. Accumulation of sugars during exposure to cold temperatures has been linked to the direct effect of low temperatures on enzyme activity involved in carbohydrate metabolism in invertebrates [66].

Seemingly, sugars also accumulate in plants exposed to temperature variation [12,67]. A recent study found that natural changes in light and varying temperature have profound impact on diel primary metabolism of *A. thaliana* compared to stable climate conditions with a constant temperature and sinusoidal simulations of light intensity [12]. Thus, most metabolites increased in the daytime and declined during the night in both the constant and variable thermal regimes, reflecting the build-up of reserves in the light and their consumption in the dark. However, the levels of sugars and starch were higher at dawn in the naturally variable regime and this pattern was associated with slow carbon utilization at night due to cold temperatures. In the variable temperature regime, the rise in amino acids was also smaller than in the stable temperature regime. This may partly be a result of decreased metabolic connectivity, which affected amino acids in particular. Whether the increased heat tolerance found for all species in the current study is a direct impact of the osmoprotective effects of sugars or an indirect effect of the accumulation of energy-yielding molecules that are readily available for other protective mechanisms remains unclear.

In *O. cincta* an increase in 3-hydroxybutyric acid and hydroxyphenyl ethanol was found in response to fluctuating temperatures (fig. 4A). The first is a ketone that is produced from fatty acid metabolism when blood glucose levels are low [68]. This could be an indication of increased energy demand, potentially because of increased metabolic rate. The latter is a phenolic compound which is synthesized by some plants and microorganisms, including fungi, bacteria, and algae species which constitute the food items for collembolans [69]. Phenolic compounds function as protective agents against herbivore and pathogen attack in plants or act as signaling molecules that attract symbionts. Additionally, phenolic compounds that are ingested via foods possibly act as antioxidants in human cells. Antioxidants protect cells from oxidative stress that results from accumulation of reactive oxygen species (ROS) [70]. High and low temperatures lead to cellular changes that induce ROS accumulation [71] and thus, the increased levels of hydroxyphenyl ethanol found in collembolans (Fig. 4) might help alleviating cells from oxidative stress. The elevated levels of this metabolite presumably come from increased food consumption, maybe as a response to increased energy demand. However, these are merely speculations and need further testing to make any conclusions.

Collectively, results of the present study show a beneficial effect of thermal fluctuations on heat tolerance in a plant species, and in two taxonomically distinct invertebrate species. *A. thaliana* had greater capacity to change its heat tolerance in response to thermal fluctuations in comparison to the two investigated invertebrate species and the underlying metabolic response was likewise stronger in *A. thaliana*. The physiological mechanisms of adaptation were comprised of metabolites involved in primary metabolism in both *A. thaliana* and *D. melanogaster*, but showed a markedly different metabolite response in *O. cincta* involving fatty acid metabolism and phenolic compounds.

## Acknowledgements

The authors thank Anders Pedersen at the Swedish NMR Center at the University of Gothenburg for help with sample preparation and experimental setup and for access to the 800 MHz spectrometer.

## Authors’ contributions

M.P., S. B., T.N.K, and N.K.N conceived the ideas, designed the methodology, and collected the data. A.M. processed samples for metabolomics analysis, and N.K.N and A.M. analysed the data. T.N.K. and N.K.N led the writing of the manuscript and all authors contributed to the drafts and gave final approval for publication.

